# Colony volatiles and substrate-borne vibrations entrain circadian rhythms and are potential mediators of social synchronization in honey bee colonies

**DOI:** 10.1101/850891

**Authors:** Oliver Siehler, Guy Bloch

## Abstract

Internal circadian clocks organize animal behavior and physiology and are entrained by ecologically-relevant external time-givers such as light and temperature cycles. In the highly social honey bee, social time-givers are important and can override photic entrainment, but the cues mediating social synchronization are unknown. Here we tested whether substrate-borne vibrations and hive volatiles can mediate social synchronization in honey bees. We first placed newly-emerged worker bees on the same or on a different substrate on which we placed cages with foragers entrained to ambient day- night cycles, while minimizing transfer of volatiles between cages. In the second experiment, we exposed young bees to constant airflow coming from either a free-foraging colony or a similar size control hive containing only empty combs, while minimizing transfer of substrate-borne vibrations between cages. After five days, we individually isolated each focal bee in an individual cage in an environmental chamber, and monitored locomotor activity. We repeated each experiment five times, each trail with bees from a different source colony, monitoring a total of more than 1000 bees representing diverse genotypes. We found that bees placed on the same substrate as foragers showed a stronger phase coherence; and in 3 of 5 trials their phase was more similar to that of foragers, compared to bees placed on a different substrate. In the second experiment, bees exposed to air from a colony showed a stronger phase coherence, and in 4 out of 5 trial their phase was more similar to that of foragers, compared to control bees exposed to air from an empty hive. These findings lend credence to the hypothesis that surrogates of activity such as substrate-borne vibrations, and volatile cues entrain circadian rhythms in natural free-foraging honey bee colonies.

## Introduction

Circadian clocks are endogenous pacemakers generating rhythms of about 24 hrs that organize animal physiology and behavior. The ubiquity of circadian rhythms in animals is consistent with the notion that circadian clocks are functionally significant because they allow animals to adjust their physiology and behavior to predicted changes in their environment (Dunlap et al., 2004; Yerushalmi and Green, 2009; Helm et al., 2017; Kumar, 2017). These internal clocks are entrained by ecologically-relevant environmental cues (known as “Zeitgebers” or “time-givers”) that are detected by sensory systems, converted to neuronal information, and transmitted to set the phase of central pacemakers. Output pathways carry temporal signals from the entrained clock to various downstream processes (Dunlap et al., 2004).

Most studies on the entrainment of circadian clocks have focused on photic entrainment which is considered evolutionary ancient and the most important time-giver to circadian clocks. In addition to light, the sun also produces daily fluctuations in ambient temperature that can effectively entrain the circadian clocks of many organisms (Sharma and Chandrashekaran, 2005; Albrecht, 2012). Non-photic, non-thermal, time-givers have received little attention. However, recent studies have highlighted the importance of entrainment by feeding time that may act on clock in tissues other the central light entrained clock (e.g., the SCN in mammals; Mendoza, 2007).

A somewhat more controversial line of research relates to the importance of social entrainment. Although observations consistent with entrainment by social interactions have been reported for diverse animal species, there is no clear relationship between the level of sociality and the efficacy of social entrainment (reviewed in Favreau et al., 2009; Castillo-Ruiz et al., 2012; Eban-Rothschild and Bloch, 2012b; Bloch et al., 2013). Some social animals such as Mongolian gerbils, sugar gliders, and common marmosets could not be effectively entrained by even intensive social interactions such as contacts with receptive females or aggressive opponents. On the other hand, social influences on activity rhythms have been reported for species which are not considered social, such as the fruit fly *Drosophila melanogaster* (Levine et al., 2002; Lone and Sharma, 2011; Bloch et al., 2013; Castillo-Ruiz et al., 2012). An additional difficulty is that no neuronal or molecular pathway transmitting social stimuli to the core clock system has been described to date. Rather, it has been suggested that social influences on the mammalian clock are mediated by changes in arousal state, by learning processes associating circadian time with specific actions, or by social gating of the time or pattern of exposure to photic time-givers (Reebs, 1989; Amir and Stewart, 1996; Mistlberger et al., 2004). The best evidence for social entrainment is found in cavity-dwelling social animals such as bats, ants, and bees in which at least some individuals do not experience ambient conditions directly, but rather rely on information received from group mates that forage outside (Regal and Connolly, 1980; Favreau et al., 2009; Eban-Rothschild and Bloch 2012b; Bloch et al., 2013; Mildner and Roces, 2017). In social insects it is thought that social synchronization of activity time is essential for temporal coordination and efficient colony-level performance. For example, honey bee nectar receivers and foragers need to act in a concerted manner and this can be achieved by means of social synchronization (reviewed in Sharma, 2003; Bloch et al. 2013; Eban-Rothschild and Bloch, 2012b).

We study the highly social (“eusocial”) honey bee (*Apis mellifera*), that has been relatively well-studied both in terms of social behavior and circadian rhythms. The organization of work in honey bee colonies relates to the age of worker bees. Young workers typically stay in the constantly dark and tightly thermoregulated cavity of the hive in which they typically care for (“nurse”) the brood. Older workers typically forage outside the hive for food and other resources (reviewed in Robinson, 1992). Foragers are diurnal and show a sleep-like behavior at night, whereas nurses typically tend brood around-the-clock (Eban-Rothschild and Bloch, 2012a; Bloch, 2010). Although around-the-clock active nurses typically do not show overt circadian rhythms in locomotor activity or whole brain clock-gene transcript abundance, there is evidence that they do have functional and entrainable circadian clocks. Firstly, nurses that are removed from the hive and isolated individually in a cage, rapidly switch to activity and brain gene expression with robust circadian rhythms that are in phase with ambient day-night cycles (Shemesh et. al., 2007; 2010). Secondly, the expression levels of more than 150 brain transcripts vary with a 24 h cycle (Rodriguez-Zas et al., 2012). Thirdly, recent immunocytochemical studies show that the levels of the clock protein *amPeriod* and the circadian neuropeptide *Pigment Dispersal Factor* (PDF) cycle with a similar phase and amplitude in brain circadian neurons of foragers that show, and nurses that do not show, circadian rhythms in locomotor activity (Fuchikawa et al., 2017; Beer et al., 2018). Here we ask what are the time-givers that entrain the clocks of nurses that dwell inside the constantly dark and tightly thermoregulated hive?

There is good evidence that the clocks of nurses are entrained by social time givers (reviewed in Bloch, 2010; Eban-Rothschild and Bloch, 2012b; Bloch et al., 2013). Moreover, Fuchikawa et al., (2016) showed that nest bees as young as two days of age can be socially entrained to the colony cycle. Entrainment was effective for caged bees deprived of access to the hive entrance and direct contact with other individuals in the colony. Thus, social entrainment cannot be explained by gating the time of exposure to light. Remarkably, social entrainment was potent enough to override photic entrainment, because forager activity stably entrained the clock even for nest bees confined to the inner dark nest cavity and experiencing conflicting photic and social environmental cycles. This evidence for the power of social entrainment highlights the need to know what are the signals and interactions mediating social entrainment in honey bees.

We hypothesize that surrogates of forager activity, such as humidity, CO_2_ levels, volatile pheromones or comb vibrations, act to synchronize circadian rhythms of individual bees and entrain the colony to a common phase (Bloch et al., 2013). Here we focused on two important surrogates of bee activity, substrate-borne vibrations and hive volatiles. Honey bees, as well as other social insects, use vibrations for intracolony communication (reviewed Hunt and Richard, 2013). Volatile pheromones and other hive odours are also well established for their pivotal and diverse communication functions in insect colonies (Wilson, 1971; Conte and Hefetz, 2008; Leonhardt et al., 2016). The hypothesis that chemical cues affect social synchronization was supported by a study showing that allowing airflow between small groups of bees facilitates their synchronization to a common circadian phase (Moritz and Kryger, 1994). There is also evidence that hive CO_2_ -levels vary over the day (Ohashi et al., 2009; Murphy et al., 2015; Edward-Murphy et al., 2016), and in other insects there is evidence consistent with the premise that CO_2_ can entrain circadian rhythms (Nicolas and Sillans,1989).

To test the hypothesis that substrate-borne vibrations generated by forager activity synchronize circadian rhythms in locomotor activity we analyzed the circadian phase of newly-emerged bees placed on the same or on different substrate as foragers. To test the second hypothesis that volatile cues from a free-foraging colony can mediate social synchronization, we exposed newly-emerged bees to a constant air flow coming out from either a hive housing a free foraging colony or a control empty hive. Our findings show that both vibrations generated by foragers activity and volatile cues from a free-foraging colony can entrain circadian rhythms in locomotor activity in young worker bees. These findings lend credence to the hypothesis that surrogates of forager activity mediate social synchronization in honey bees.

## Material and Methods

### Bees

Honey bees were maintained according to standard beekeeping techniques at the Bee Research Facility at the Edmond J. Safra campus of the Hebrew University of Jerusalem, Givat-Ram, Jerusalem, Israel. The honey bees in our apiary are derived from a mixture of subspecies typical to Israel. To obtain newly-emerged bees, we removed honeycomb frames with emerging worker pupae, brushed-off all adult bees, and immediately transferred each frame into a separate lightproof container. We placed the frames in an incubator (33°C ±1 °C, 60%RH ±5%) for the bees to emerge. The emerging bees were collected from the comb within two hours post emergence under dim red light (DD; using Edison Federal EFEE 1AE1 Deep Red LED; mean wavelength = 660 nm, maximum and minimum wavelengths = 670 and 650, respectively) light to avoid influences of light on the circadian system (e.g., photic entrainment).

### General procedure

Fig. 1A summarizes the general experimental outline. On Day-1, we collected newly-emerged worker bees and transferred them to Libfield wooden cages (12 × 8 × 4.5cm) with glass covers in which they were exposed to the tested environmental cues. On Day-6, we collected samples of bees experiencing the various treatments, as well as foragers that served as a control group. We isolated each focal bee in a monitoring cage and monitored locomotor activity for at least seven successive days under dim red light and constant lab conditions (Fig. 1A). We used the locomotor activity data to determine the circadian phase of each bee and assess social synchronization among bees subjected to the same treatment (e.g., colony odors or vibrations).

**Figure 1.**
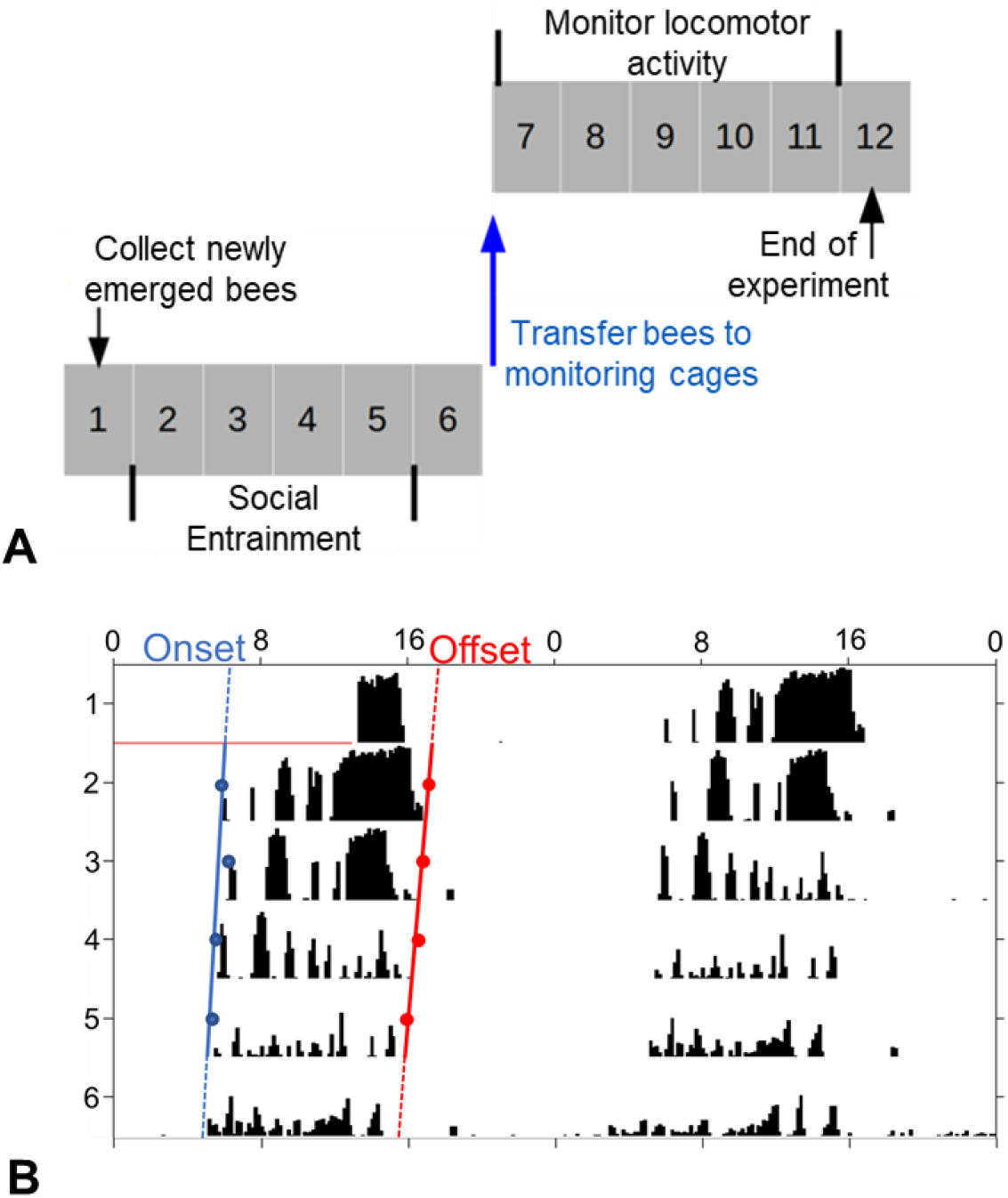
**(A) General experimental outline.** The numbers depict the days of the experiment and the grey filling indicate that the bees were kept under dim red light. On Day-1 (1), we collected the bees and transferred them to cages in the lab, in which they were exposed to the tested signals. On Day-6, each focal bee was collected and transferred to an individual monitoring cage in which her locomotor activity was automatically monitored for at least seven successive days. **(B) Representative double-plotted actogram of an individually isolated bee.** The y-axis shows days in the monitoring cage, and the X-axis the time of day, double plotted for easier visual detection of rhythms. The height of the black bars within each day corresponds to the level of locomotor activity in a 10 min. bin, determined by distance moved in pixels (translated to millimeters) [mm] in 1-min. The blue and red dots show the estimated times for the onset and offset of activity for each day, respectively. Linear regression models are fitted to these points and the phase is determined based on the extrapolation of the regression lines on Day-1.

### Monitoring and analyzing locomotor activity

We placed each bee individually in a monitoring cage made of a modified Petri dish (diameter = 90mm) provisioned with *ad libitum* sugar syrup (50% w/w) and pollen. The monitoring cages with the bees were placed in a tightly regulated environmental chamber (29 ±°1C, 55 ±5% RH). The chamber was illuminated with dim red light (Edison Federal EFEF 1AE1 Far (Cherry) Red LED; mean wavelength = 740 nm, maximum and minimum wavelengths were 750 and 730, respectively). Locomotor activity (measured as number of pixels traveled over a time unite on the camera field of view and transformed to millimeters) was recorded automatically at a frequency of 1 Hz with the ClockLab data acquisition system (Actimetrics Inc., Evanston, IL, USA). Our monitoring system is composed of four infra-red light-sensitive black and white cameras (Panasonic WV-BP334, 0.08 lux CCD video cameras or Sentech STC-MB33USB mini USB video cameras with Computer TZ32910CS-IR lenses) and a high-quality monochrome image acquisition board (IMAQ 1409, National Instruments) or a National Instruments USB-6501 interface. Each camera records the activity of 30 cages placed on a single tray. Thus, in each trial we could monitor up to 116 bees; additional 4 empty cages, one on each tray, were used as a control recording background noise.

For the analyses of circadian rhythms, we used the ClockLab circadian analyses software package (Actimetrics, USA). We used the χ^2^ periodogram analysis with 10- minute bins to generate actograms for the analyses of circadian rhythms. As a proxy for the strength of circadian rhythms we used the *Power* which was calculated as the height of the periodogram peak above the P = 0.01 significance threshold line (for more details see Yerushalmi et al., 2006). For each day we calculated the onset and offset of the daily bout of activity. The precise timing of the onset or offset were defined as at least three consecutive 10-min bins each with activity reaching at least 10% of the maximal activity per bin during this day. In addition, there was a period of at least 5 hrs of reduced activity between the offset and the following onset (as described in Fuchikawa et al., 2016). We used the ClockLab software to fit linear regression models passing through the onset or the offset points of at least four consecutive days. The extrapolations of these regression lines on the first day in which the bees were transferred to the monitoring system were used for estimating the timing of onset and offset of activity (Fig. 1B). We included in our analyses only bees with statistically significant circadian rhythms (χ^2^ periodogram analysis; P < 0.01, period between 20-28 hrs.) for which we could unambiguously determine the onset/offset of activity.

We used the Oriana circular statistics software package (KCS, USA) to determine the degree of synchronization and the phase coherence among bees within each treatment group. We used the time of onset, offset, and median (which was defined as the midpoint between the times of onset and offset) as indices for the phase. The Rayleigh test was used to determine if phase synchronization is significantly different from random distribution, and the mean length of the Rayleigh vector was used as an index for phase coherence. We used the Watson Williams F-test to compare the phases of two groups.

#### The influence of substrate-borne vibrations on social synchronization of circadian rhythms in locomotor activity

This experiment was designed to examine the hypothesis that substrate-borne vibrations can mediate social synchronization among honey bee workers. The experimental design is summarized in Fig. 2. On the first day, we caged groups of ~30 callow (i.e., newly emerged) bees or foragers in Libfield wooden cages (12 × 8 × 4.5cm) with glass covers, provisioned each cage with ad libitum sugar syrup (50% w/w and pollen, collected by honey bees). Foragers were collected at the hive entrance and identified as bees returning to the hive with pollen loads conspicuously attached to their hind legs. They were collected on the same day and from the same source colony as the callow bees. The foragers were then entrained to a light-dark cycle that was 5 hours advanced relative to the phase of the colony in the first trial, and to the natural day-night cycle in all other trials. We placed the cages containing groups of callow bees either on the same substrate (a canvas frame placed on a table) as the foragers (“*Same Substrate”*) or on a different substrate (*“Different Substrate”*). In each trial we used two cages for each treatment (see Fig. 2). We placed all the cages in the same environmental chamber (30 °C ±1°C, 55% RH ±5%, illuminated with dim red Edison Federal EFEE 1AE1 Deep Red LED lights, mean wavelength = 660 nm, maximum and minimum wavelengths = 670 and 650, respectively). The distance between the cages of foragers to those with callows on the *Same Substrate-* and *Different Substrate*- was identical (Figure 2). We sucked out volatiles (that may mediate social entrainment) using an exhaust air unit with its opeining (diameter 10cm) placed above the foragers cages. We repeated this experiment five times between May to August 2017, each trail with bees from a different source colony. The survival rate was 45% (range 30% - 70%) for foragers and 87% (range 61% - 100%) for young bees. Such differences in survival are common because foragers are older and foraging activity is a physically demanding activity.

**Figure 2:**
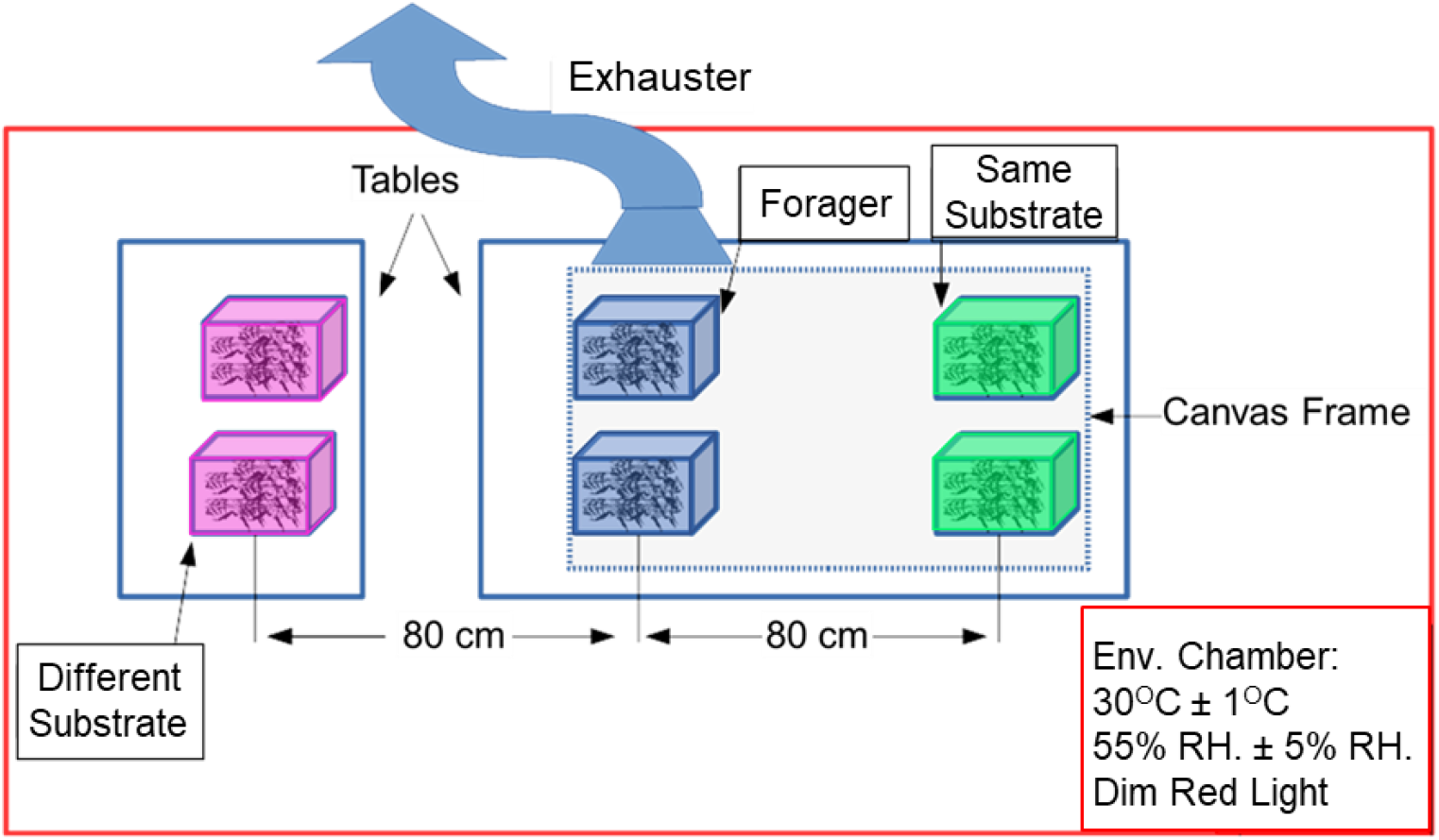
Setup for the experiment testing whether substrate-borne vibrations entrain circadian rhythms in locomotor activity. We placed newly emerged bees in two small cages (30 bees/cage) on the same (*‘Same Substrate’*) or on a different substrate (*‘Different Substrate’*) on which we placed two cages with foragers. We sucked out the air above the foragers to prevent the spread of forager emitted odorants that could influence circadian rhythms of young bees.

#### The influence of colony volatiles on social synchronization of circadian rhythms in locomotor activity

To test the hypothesis that volatile chemicals mediate social synchronization of honey bee workers, we exposed callow bees to volatiles pulled out from a regular hive housing a free-foraging colony (“*Colony”*), or from a control empty hive (*“Empty Hive”*). The empty hive was of a similar size and contained the same number of empty combs. To keep the temperature in the *Empty Hive* constant at a similar level as in the *Colony*, we heated it with a “Reptile-heating mat” (20W- 16.5×11 in). Both hives were prepared with a similar hole in the back board where an exhauster tube (diameter 25mm) was installed (Figure 3).

**Figure 3.**
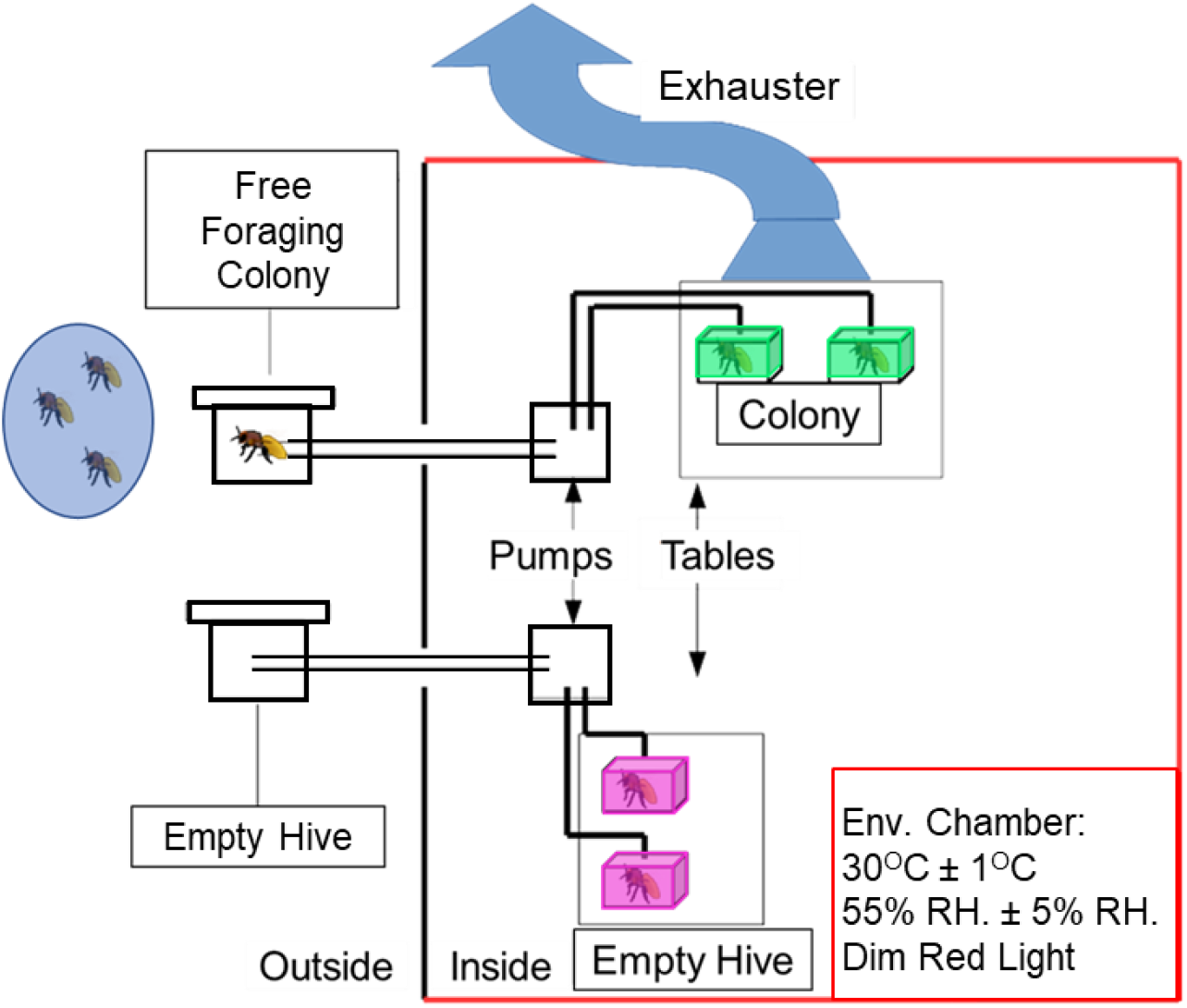
Setup for the experiment testing whether hive volatiles entrain circadian rhythms in locomotor activity. Cages, each with 30 callow bees, were exposed to a constant air flow sucked out from either a hive housing a free-foraging colony (*‘Colony’*) or a similar control empty hive containing the same number of empty combs (*‘Empty Hive’*). We placed all the cages on vibration-absorbing bases to minimize possible transfer of vibrations. We sucked the air above the *Colony-group* to prevent unintended release of volatiles that could affect the behavior of other bees.

We pumped the air out of the hive boxes using two aquarium pumps (Atman, Aquarium Air Pump AS-1063). The air from each hive was flowed into two Libfield cages, each containing 30 callow bees (Fig. 3). The callow bees in these cages were provisioned with ad libitum sugar-syrup (50% w/w) and pollen. We placed the cages on vibration-absorbing bases to minimize possible transfer of vibrations. We placed a tube (diameter 10cm) from the exhaust-air unit above the cages of the *Colony-group* in order to prevent unintended transfer of volatiles to the Control cages. The pump and the cages were housed inside a tightly regulated environmental chamber (30 ±1oC, 55 ±5% RH) constantly illuminated with dim red light (as in the first experiment above). The populated and empty hives were placed outside the building. The colony in the populated hive was self-sustained (Fig. 3). We repeated this experiment five times between September to November 2017, each trail with bees from a different source colony. The average survival rate was 79% for foragers (range 70 - 100%) and 67% for the young bees (51 - 84%).

## Results

### The influence of substrate-borne vibrations on social synchronization of circadian rhythms in locomotor activity

We repeated this experiment five times, each trail with bees from a different source colony, and altogether monitored 454 bees. The experimental groups differed in the strength of circadian rhythms. The foragers had stronger rhythms (Power = 222.76 ±160.09 SD) compared to the two groups of young bees, that were overall similar (“*Same Substrate”:* 128.89 ±119.81, and *“Different Substrate”:* 122.52 ±115.04). The proportion of bees showing statistically significant circadian rhythms was 91% (range 70-100%) for *Foragers,* 71% (range 58-85%) for young bees on the *Same Substrate,* and 68% (range 50-70%) for the *Different Substrate-group* (data not shown).

The phase of *Same Substrate* bees was overall more similar to that of the foragers compared to that of the *Different Substrate* treatment. In 4 out of 5 trails, the phase of the *Different Substrate-group* was significantly different from that of the foragers (Watson Williams F test < 0.05; Fig. 4A). The young bees on the same substrate also showed stronger phase coherence (longer Rayleigh vectors) compared to those placed on a substrate different from that of the foragers (Fig. 4B; Kruskal-Wallis test p = 0.007; Steel-Dwass-Critchlow-Fligner *post hoc* test: *Forager – Same Substrate p* = 0.417; *Forager - Different Substrate, p* = 0.024; *Same Substrate – Different Substrate* = 0.116). We further compared the phase-difference (differences in minutes between the mean vectors) relative to the foragers. In all five trials, the phase difference between the *Foragers-* and the *Different Substrate-group* was larger than compared to the *Same Substrate-group* (Binomial test, n = 5, p = 0.031) (Fig. 4C).

**Figure 4.**
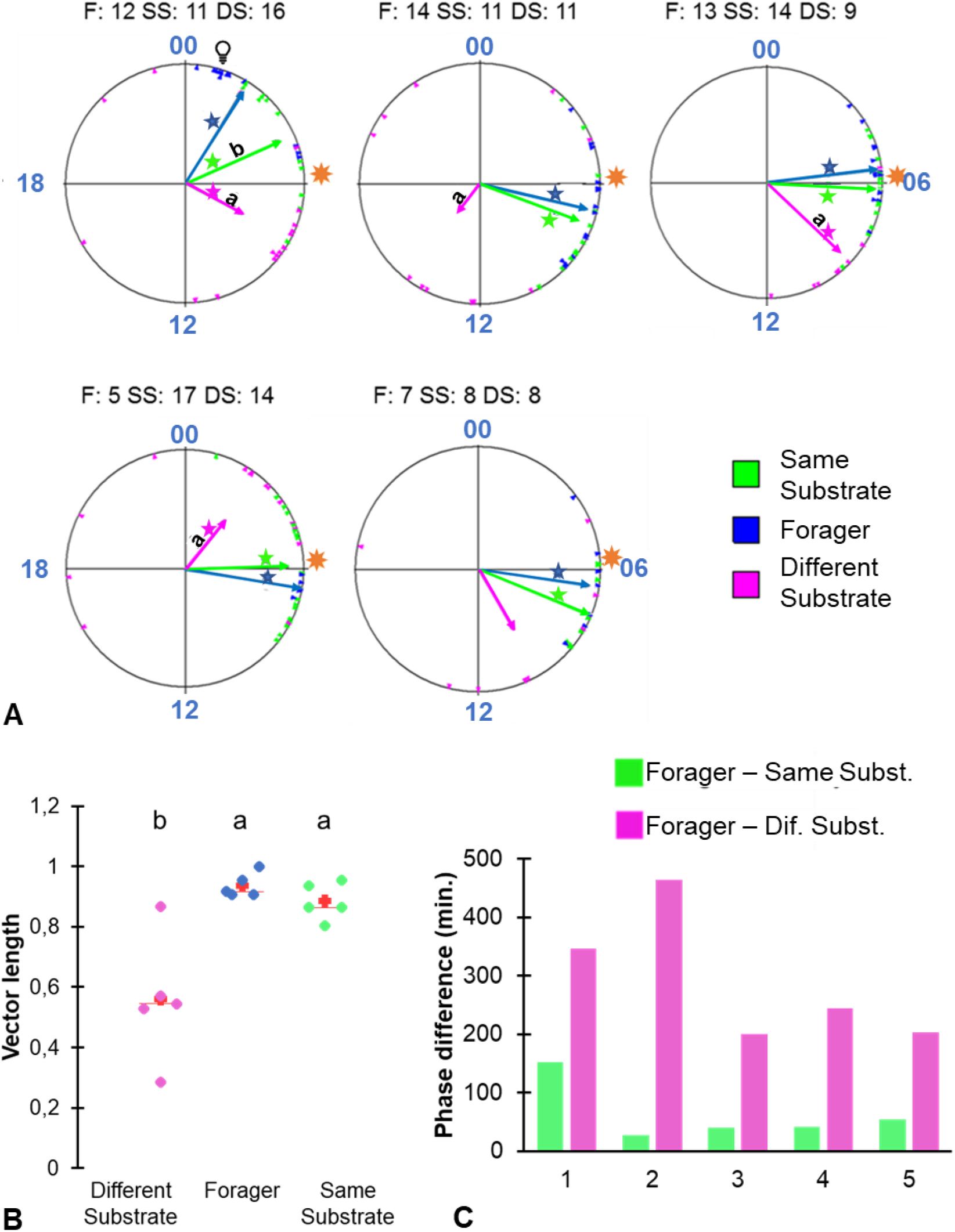
Substrate-borne vibrations entrain circadian rhythms in honey bee workers. **(A)** Circular plots comparing the phases of bees from the different treatment groups. Each plot summarizes a different trial. Sample sizes (number of bees with a clear onset) are shown above each plot. F= foragers; SS = callows on the same substrate; DS = callows on a different substrate. The time of day is depicted on the circular plot perimeter. Each triangle depicts the onset of an individual bee. The vectors point to the average onset time, and their length corresponds to the degree of phase coherence. Asterisks indicate the p-value obtained from Rayleigh test is significant (*- α < 0.05). Vectors marked with different letters differ in phase in a Watson-Williams F-test. The foragers were entrained to a phase 5 hours advanced (illustrated by the light bulb symbol) relative to the phase of the colony (illustrated by the sun symbol) in the first trial, and to the natural sunrise in all other trials (symbol). **(B)** A Scatter plot summary of the degree of synchronization (length of the Rayleigh vector) across the different treatments. The red crosses correspond to the means and the central horizontal bars to the medians. Treatments marked with different letters are significantly different in a Kruskal-Wallis Test (P<0.05) followed by Steel-Dwass-Critchlow-Fligner *post hoc* test. **(C)** The phase-difference between each of the two groups of young bees relative to the foragers.

### The influence of hive volatiles on social synchronization of circadian rhythms in locomotor activity

We performed five repetitions of this experiment, each with bees from a different source colony. In total, we monitored the locomotor activity of 580 bees. As in the first experiment, the foragers had stronger circadian rhythms (mean Power = 234.43 ±119.48 SD) than the two groups of young bees which were overall similar (*Colony-* 166.93 ±115.44; *Empty hive-group-* 158.94 ±127.31). Almost all the foragers (98%; range 95-100%) showed statistically significant circadian rhythms, whereas only 75% (range 62-89%) in the *Colony-* and 70% (range 58-84%) in the *Empty hive-group* showed significant circadian rhythms (Data not shown).

The phase of the young bees that were exposed to air from a hive housing a colony (“*Colony*”) was more similar to that of the foragers (collected from the same colony from which we sucked the air) compared to that of the callow bees exposed to air sucked from a similar hive with no bees. In 3 out of 5 trails, the phase of the *Empty hive bees* was significantly different from that of the *foragers* (Watson Williams F-Test, p < 0.05; Fig. 5A). Bees from the *Colony* treatment showed stronger phase coherence (longer Rayleigh vector) compared to the Empty hive treatment (Kruskal-Wallis test, p < 0.0001; Steel-Dwass-Critchlow-Fligner *post hoc* test: *Forager-Colony*, p = 0.043; *Forager-Empty hive*, p = 0.024; *Colony-Empty hive* p = 0.043) (Fig. 5B). In a pooled analysis across all five trials, the phase-difference relative to the *Foragers*, was similar for the two groups of young bees (Binomial test p = 0.31; Figure 5C). It is notable however, that in the first two trials the phase difference was much larger for the bees exposed to the empty hive.

**Figure 5.**
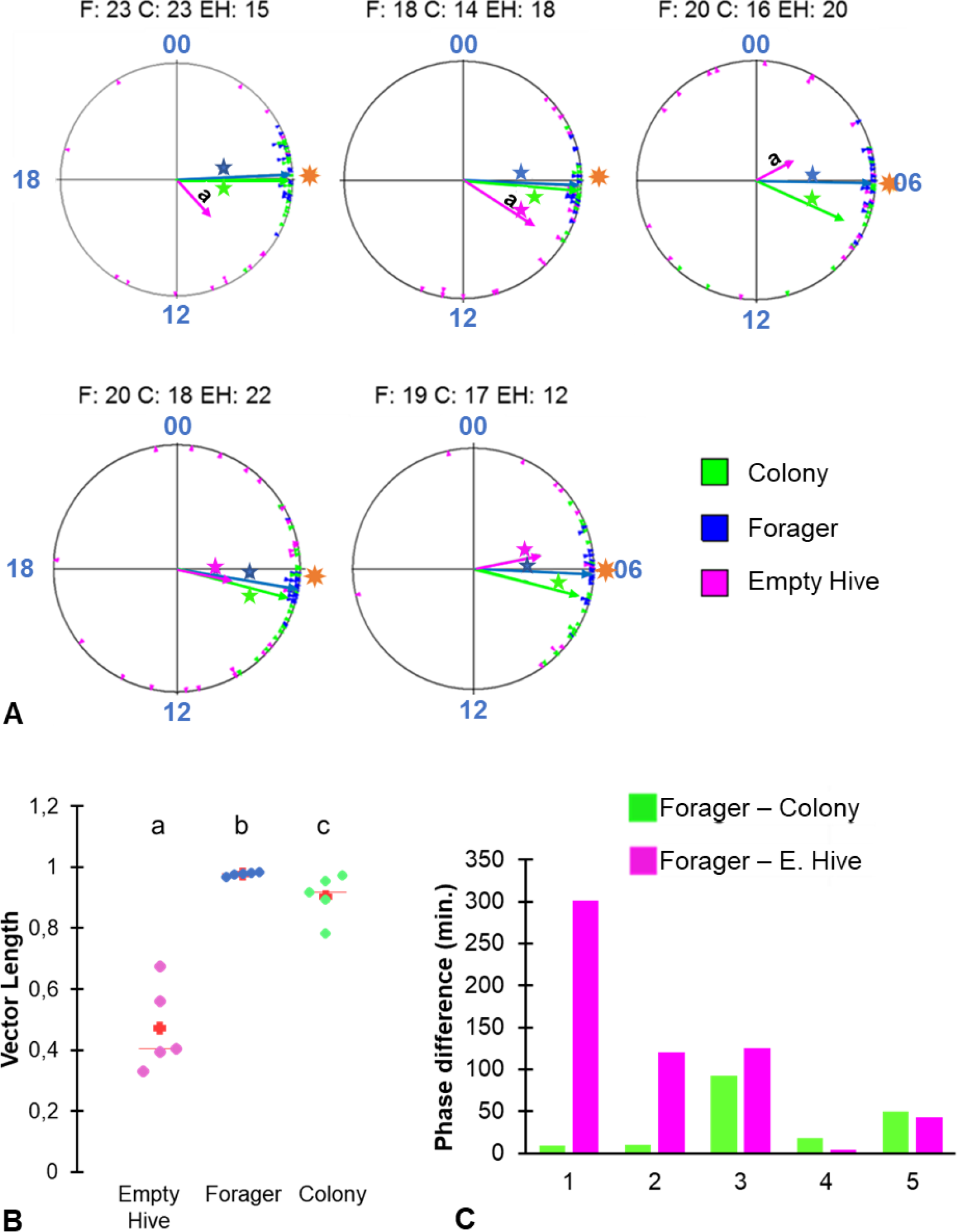
Hive volatiles mediates social synchronization: **(A)** Circular plots comparing the phases of bees from the different treatment groups. Each plot summarizes a different trial. Sample sizes are shown above each plot (number of bees with a clear onset) F= foragers; C = callows exposed to air from a colony; EH = callows exposed to air from an empty hive). The time of day is depicted on the circular plot perimeter. Each triangle depicts the onset of an individual bee. The vectors point to the average onset time, and their length corresponds to the degree of phase coherence. Asterisks indicate the p-value obtained from Rayleigh test (*- α < 0.05). Vectors marked with letters have a significantly different phase in a Watson-Williams F-test. The foragers were entrained by the natural sunrise (illustrated by the sun symbol) in all trials. **(B)** A Scatter plot summary of the degree of synchronization (length of the Rayleigh vector) across different treatments. The red crosses correspond to the means and the central horizontal bars are the medians. Treatments marked with different letters are significantly different in a Kruskal-Wallis Test (P<0.05) Steel-Dwass-Critchlow-Fligner *post hoc* test, *Forager-Colony* p = 0.043; *Forager-Empty hive* p = 0.024; *Colony-Empty hive* p = 0.043. **(C)** The phase-difference in minutes between each of the two groups of young bees relative to the foragers.

## Discussion

Our experiments show that both substrate-borne vibrations generated by forager activity, and volatile cues emitted by a free-foraging colony can entrain circadian rhythms in locomotor activity of young honey bees. These results were obtained using a robust dataset composed of more than 1000 bees that were each monitored individually. We repeated each experiment five times, each trial with bees from a different source colony. Given that bees in each colony are the offspring of a different queen and drones, our findings are therefore not limited to certain genotypes or laboratory lines. It is also important to note that given that we determined the circadian parameters for bees after removing them from the environment in which they were entrained, our measurements reflect properties of the internal clock rather than environmental (e.g. social) factors that could mask the clock effect on locomotor activity. Our experimental approach allowed us to uncouple the effects of substrate-borne vibrations and volatile colony cues. In the experiment testing the influence of substrate vibrations, we minimized the transfer of volatiles by sucking out the air above the forager cages. In the experiment testing the influence of hive volatiles, we minimized possible effects of vibrations by placing the cages on vibration-absorbing bases. The conclusive evidence for synchronization in both experiments suggest that both vibrations and volatile cues are potent enough to mediate social synchronization. Taken together, these findings lend credence to the hypothesis that vibrations and volatile olfactory cues, that are generated by active bees, mediate social synchronization among worker bees in a honey bee colony.

These findings are consistent with the hypothesis that volatile cues in the hive mediate social synchronization in honey bee colonies (Moritz and Kryger, 1994; Bloch et al., 2013; Fuchikawa et al., 2016). We show that young bees exposed to volatiles from a hive housing a free-foraging colony are better synchronized with each other and overall have a phase more similar to that of the free-foraging colony, compared with similar bees that were exposed to air coming from a similar hive with no bees (Fig. 5). These findings are consistent with earlier studies showing that young bees that were caged inside double mesh enclosures were similarly synchronized with the ambient day- night cycles as bees freely moving in the hive, indicating that direct contact is not necessary for social synchronization in honey bees (Fuchikawa et al., 2016). Similarly, Moritz and Kryger (1994), showed that the synchronization among bees in two small groups was improved when they allowed air to flow between the two groups. What are the colony volatiles mediating this effect? We propose that the two most likely factors are volatile pheromones or fluctuations in CO_2_ concentrations. The CO_2_ concentration within the hive shows a clear diurnal cycle (Murphy et al., 2015; Edward-Murphy et al., 2016; Ohashi et al., 2009). Typically, lower levels are recorded during daytime when the forager are outside the hive, and high levels at the morning and the late afternoon. These two CO_2_ peaks apparently correspond to the time when a large cohort of foragers leave the nest for their first morning foraging trips, and then the evening peak is when all (or most) of them return (Ohashi et al., 2009).

Honey bees act to homeostatically regulate their nest microenvironment. To regulate CO_2_ levels, workers stand near the hive entrance and fan with their wings ventilating the nest cavity. There is a positive correlation between CO_2_ levels inside the hive and the number of fanning bees (Seeley, 1974; reviewed in Guerenstein and Hildebrand, 2008). Thus, we assume that CO_2_ levels can entrain the circadian clock of bees by two non-mutually exclusive mechanisms: First, elevated in CO_2_ levels lead to an increase in the number of bees fanning and this activity entrains their circadian clocks. Second, fluctuation in CO_2_ levels act as a zeitgeber setting the phase of circadian clocks affecting locomotor activity. Although we are aware of only a few evidences that CO_2_ can entrain circadian rhythms in insects (Nicolas and Sillans, 1989), there is better evidence for such effects for mammals. Adamovich et al., (2019) reported that changes in carbon dioxide concentration act at the cellular level and can phase shift oscillations in clock gene expression in a cell culture. Similar cellular mechanisms may be also found in honey bee and other insects.

Honey bees use a wide variety of vibrational signals to communicate and coordinate colony-level activities. For example, both the “waggle dance” with which foragers recruit followers to a rewarding flowering patch, and the “tremble dance” with which they inhibit further recruitment, are at least partially mediated by comb vibrations (reviewed in Hrncir et al., 2005; Hunt and Richards, 2013). The hypothesis that substrate-borne vibrations can mediate social synchronization is also consistent with findings that in the fruit fly *Drosophila melanogaster* circadian rhythms of locomotor activity can be entrained by oscillations in vibration intensity (Simoni et al., 2014). We do not know yet of any study recording comb vibrations over several days in honey bee colonies. Therefore, we can only speculate that forager activity which changes substantially during the day (Moore et al., 1998; Bloch and Robinson, 2001), generate comb vibrations that are sensed by nestmates and can mediate clock entrainment.

Our experiments show that comb vibrations and colony odors can synchronize circadian rhythms in honey bee workers. These findings lend credence to the hypothesis that surrogates of worker activity mediate social synchronization of circadian rhythms in honey bee colonies. The link between worker activity and comb vibration is quite straightforward, although the repertoire in the hive is broad (Hrncir et al., 2005; Hrncir et al., 2019; Hunt and Richards, 2013) and additional work is needed for identifying the specific vibrations that entrains the clock. As for colony odors, additional information on the specific of volatile chemicals that meditate this effect is clearly needed. Changes in carbon dioxide and oxygen concentration inside the hive are influenced by worker activity and metabolism. However, given that active bees typically increase their body temp’, their level of activity may also affect the release (e.g. evaporation) of additional chemicals such as cuticular pheromones from their body surfaces may be also important. We suggest that in natural colonies, these two factors, and perhaps additional surrogates of worker activity, act together to create oscillations in the nest microenvironment. This idea is consistent with self-organization models in which the sum activity of individual workers produces oscillations in the microenvironment of the hive which in turn entrain the circadian clocks of other individuals in the colony (Moritz and Fuchs, 1998; Camazine et al., 2003).

## Acknowledgment

We thank Rafi Nir for professional beekeeping assistance. We also thank Muki Nagari, Jacob Holland and Igor de Medici for assisting with the experiments.

